# MRI-based cell tracking of OATP-expressing cell transplants by pre-labeling with Gd-EOB-DTPA

**DOI:** 10.1101/2023.08.04.552007

**Authors:** Tapas Bhattacharyya, Christiane L. Mallett, Erik M. Shapiro

## Abstract

**Purpose:** To detect cell transplants labeled with a clinical gadolinium-based contrast agent as hyperintense signals using a clinically familiar T1-weighted MRI protocol.

**Methods:** HEK293 cells were stably transduced to express human OATP1B3, a hepatic transporter that transport Gd-EOB-DTPA into cells that express the transporters, the intracellular accumulation of which cells causes signal enhancement on T1-weighted MRI. Cells were labeled in media containing Gd-EOB-DTPA for MRI evaluation and indocyanine green for cryofluorescence tomography validation. Labeled cells were injected into chicken hearts, in vitro, after which MRI and cryofluorescence tomography were performed in tandem.

**Results:** OATP1B3-expressing cells had substantially reduced T1 following labeling with Gd-EOB-DTPA in culture. Following their implantation into chicken heart, these cells were easily identified in T1-weighted MRI. Cryofluorescence tomography validated that the areas of signal enhancement in MRI overlapped with areas of indocyanine green signal, indicating that signal enhancement was due to the transplanted cells.

**Conclusion:** OATP1B3-expressing cells can be pre-labeled with Gd-EOB-DTPA prior to injection into tissue, affording the use of clinically familiar T1-weighted MRI to robustly detect cell transplants.

## Introduction

Non-invasive imaging can play a key role in clinical cell therapies by determining the precise locations of transplanted cells. MRI-based cell tracking is most useful when the high-resolution and spatial discrimination capabilities of MRI are used for localization of transplanted cells and providing soft tissue anatomical context. Traditionally, iron oxide nanoparticles are used for magnetic cell labeling with detection of labeled cells via T2/T2*-weighted MRI^1^, with sensitivities as low as single cells^2^ and even single particles^3^. However, two problems have plagued iron oxide-based MRI-based cell tracking since its inception: 1) the T2/T2*-weighted dark contrast obscures the underlying anatomy, making quantification of cell number difficult, and 2) the iron oxide nanoparticles can still create MRI contrast even if the original labeled cell is long dead. Bright contrast methods have been sparingly used for MRI-based cell tracking (excellent review in ^4^) employing Gd-chelates and Mn^2+^, accumulated intracellularly by disruptive (photoporation, transfection, etc) methods or by simple incubation, respectively. Further, ^19^F agents have been used for MRI-based cell tracking^5^. Yet, MRI-based cell tracking has not made significant clinical impact, partly due to the complications of cell labeling and detection.

Hepatic organic anion transporting polypeptides (**OATPs**) are a new category of MRI reporter proteins^6-12^. OATPs are ∼700 amino acid membrane proteins with 12 membrane-spanning helices, whose expression is conserved among vertebrates ^13^. Hepatic OATPs transport off-the-shelf, FDA-approved, clinically used, MRI contrast agents into cells ^14-15^. In clinical scenarios, following IV injection of Gd-EOB-DTPA or Gd-BOPTA, two FDA-approved hepatospecific MRI contrast agents, hepatocytes become hyperintense on T1-weighted MRI due to the intracellular accumulation of the Gd-based contrast agent. This is used clinically to detect tumors in the liver as tumors (generally) do not express OATPs and remain hypointense in relation to the bright liver ^16^. As it relates to the use of OATPs as MRI reporter proteins, OATPs with reported efficient transport of Gd-EOB-DTPA include human/primate OATP1B1 and OATP1B3 ^14^, rat OATP1A1 ^8, 11^, and rat OATP1B2 ^17^. Many other species, including mouse ^18-20^, rabbit ^21^, dog ^22-24^ and pig ^25^, exhibit hepatic accumulation of Gd-EOB-DTPA and Gd-BOPTA, as evidenced by liver MRI, so it is likely that there are other members of the hepatic OATP1B family (dog OATP1B4, e.g.) that transport these agents as well.

To date, the use of hepatic OATPs as reporter proteins has relied on the IV injection of Gd-EOB-DTPA or Gd-BOPTA to accumulate in cells post-transplant ^6-12^. We hypothesized that OATP1B3-expressing cells can be pre-labeled with Gd-EOB-DTPA prior to injection affording the use of clinically familiar T1-weighted MRI to robustly detect cells immediately post-transplantation, similarly to other cell labeling paradigms. This straightforward approach to labeling and MRI detection may facilitate the incorporation of MRI-based cell tracking in clinical trials and cell therapies.

## Materials and methods

Lentivirus (Vectorbuilder) was used for stable transduction of human OATP1B3 in mammalian cells. HEK293 cells were infected at MOI 10:1 and underwent 3-week antibiotic selection to create stably expressing cells. Stable transgene expression was verified by rtPCR.

For cell transplantation studies, cells were grown to 80% confluency and then labeled in cell culture media with 5.0 mM Gd-EOB-DTPA and 2 μg/ml **ICG** for 1.5 hours, after which cells were washed. To validate labeling, an aliquot of cells was pelleted and T1 was measured at 7.0T (Bruker Biospec). ICG cellular uptake was also validated using fluorescence microscopy (Biotek Cytation3). Non-transduced cells were also similarly incubated and imaged to serve as control cells. For transplantation, 5x10^6^ dual labeled (Gd-EB-DTPA and ICG) OATP1B3-expressing cells were pelleted and resuspended in 50 μl PBS. Food-grade chicken hearts were used as a model. 1x10^6^ cells in 10 μl PBS was slowly injected free-hand into the left ventricular wall of chicken heart (n=5) using a Hamilton syringe and 22 gauge needle. Injected hearts were immediately imaged at 7.0T by 3D T1-weighted gradient echo MRI with parameters: TR/TE: 30/2.5, 1 average, FA 60°, resolution 250 um, FOV 40x30x30 mm, 8 min acquisition.

MR images were analyzed in PMOD. 3D volumes of interest (**VOIs**) were drawn in non-injected heart region, region outside the heart (noise) and the hyperintense area from the injected cells. Contrast-to-noise ratio (**CNR**) was calculated.

After MRI, hearts were processed for cryofluorescence tomography (**CFT**) (Xerra, EMIT) to validate the location of bright MRI signals. Hearts were frozen over dry ice then embedded vertically in OCT in a 7.5x9.5x5 cm mold. White light and fluorescence images (excitation 780 nm, emission 835 nm) were acquired with 30 μm in-plane resolution and 50 μm slice thickness. Image stacks were combined in PMOD and maximum intensity projections of the fluorescence images were computed.

## Results

Stable OATP1B3 expression in HEK293 cells was confirmed by qPCR and phenotypically by uptake of Gd-EOB-DTPA and ICG (Figure 1). T1 time for OATP1B3-expressing cells were 52 ms following incubation, while non-expressing cells was ∼1500 ms. ICG uptake into cells was verified microscopically. Following injection of Gd-EOB-DTPA and ICG labeled cells into the hearts, these cells appeared as hyperintense signals on T1-weighted MRI. The calculated volume of cells generated from MRI VOIs was 6.7 +/- 2.3 μl, close to the intended 10 μl injected volume, with CNR 38.4 +/- 15.4. CFT validated near identical overlap of injected cells, verifying that the bright MRI signal was generated from the ICG-labeled transplanted cells (Figures 2 and 3).

**Figure 1:**
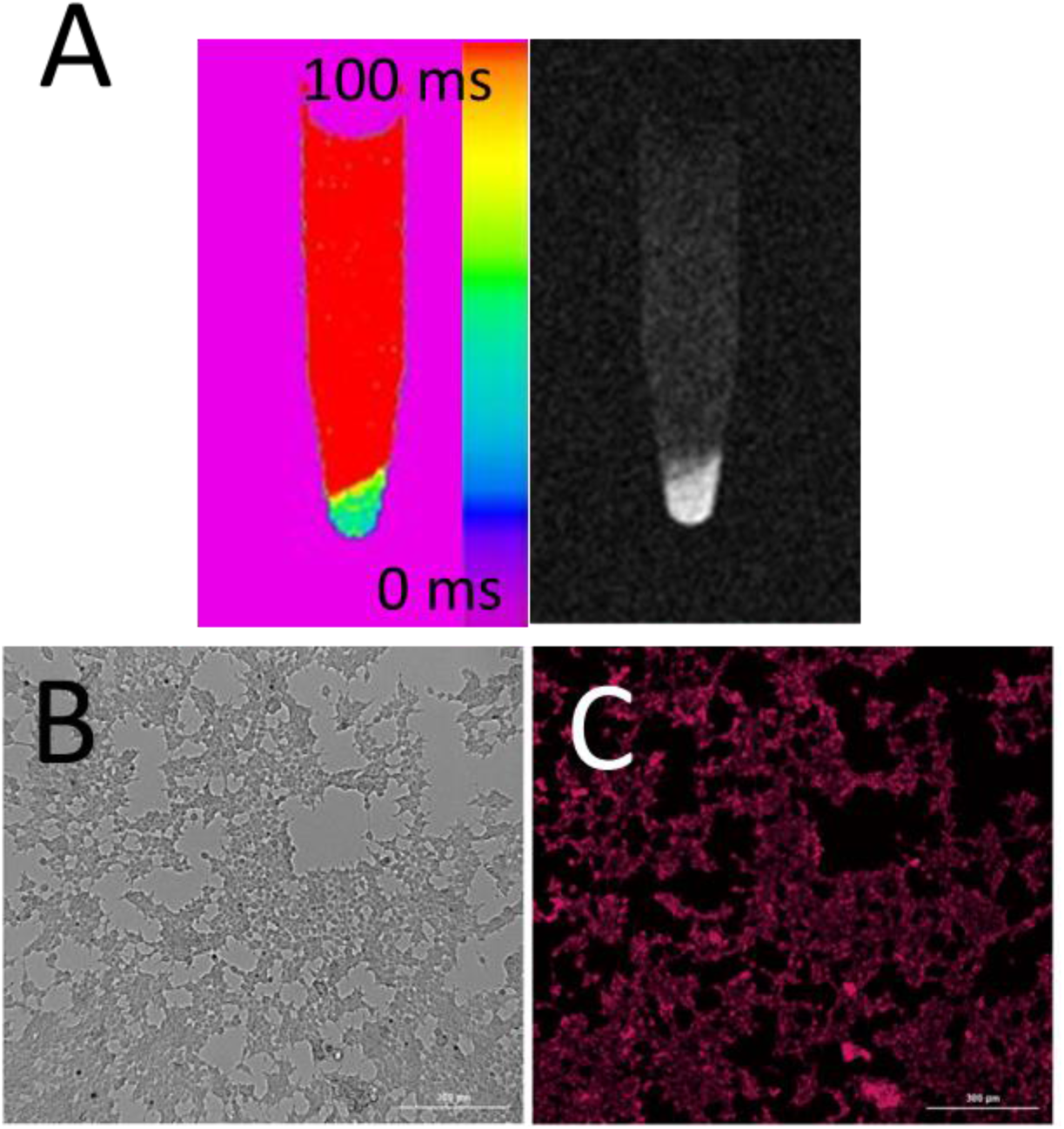
**A)** T_1_ map and corresponding MRI of OATP1B3-expressing cells following incubation in 5.0 mM Gd-EOB-DTPA and 2.0 μg/ml ICG for 1.5 hours. **B**,**C)** White light (B) and NIR fluorescent image of OATP1B3-expressing cells harboring intracellular ICG following labeling.

**Figure 2:**
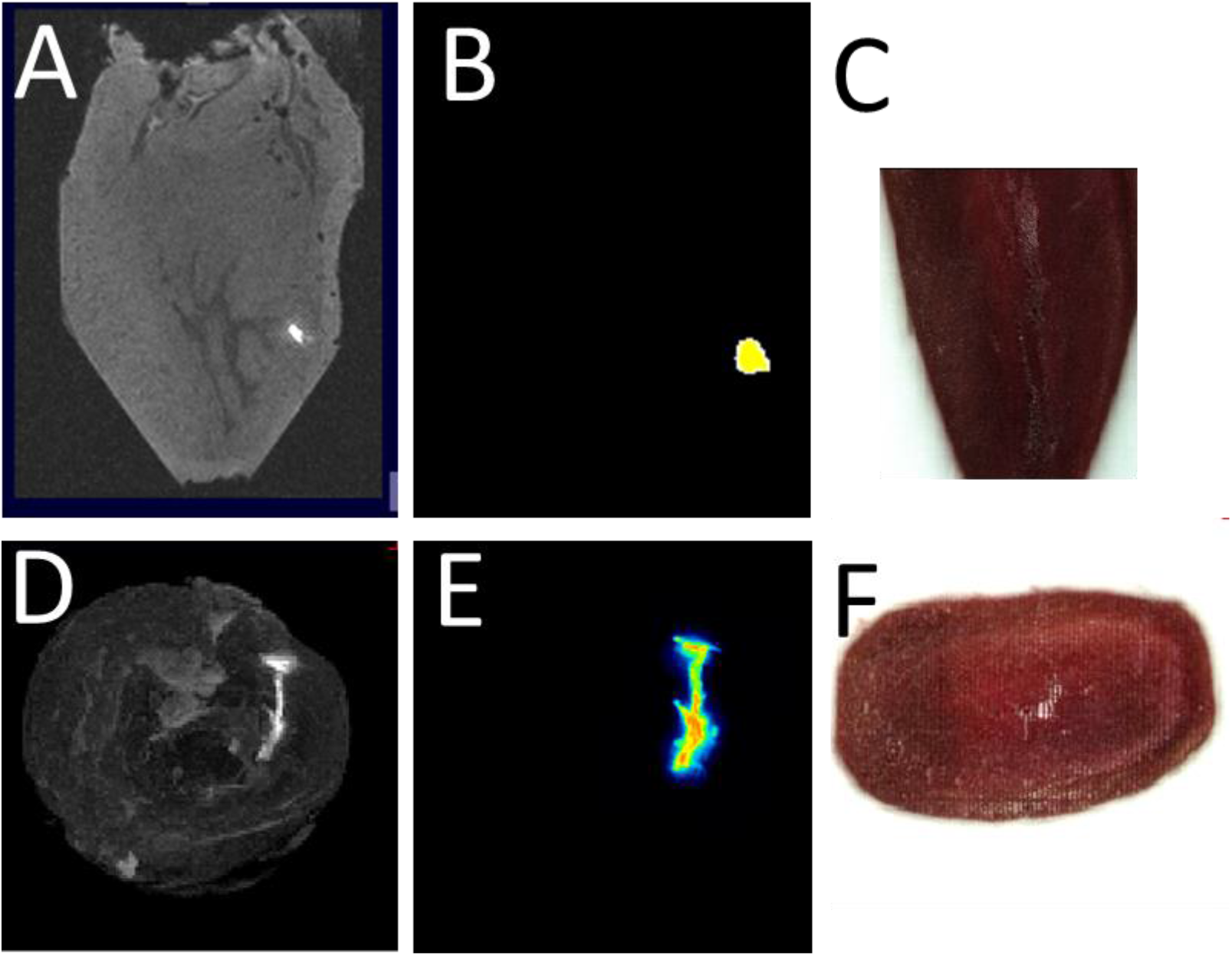
**A,B)** MRI of chicken heart with bright contrast spot from Gd-EOB-DTPA labeled OATP-expressing cells (A) confirmed by ICG NIR fluorescence from CFT image (B). **C)** White light image corresponding to area in (B). **D**,**E)** Maximum intensity projection image from MRI of entire heart showing bright contrast from Gd-EOB-DTPA labeled OATP-expressing cells (D) confirmed by ICG NIR fluorescence from CFT image (E). **F)** White light image reconstructed to correspond to area in (E).

**Figure 3:**
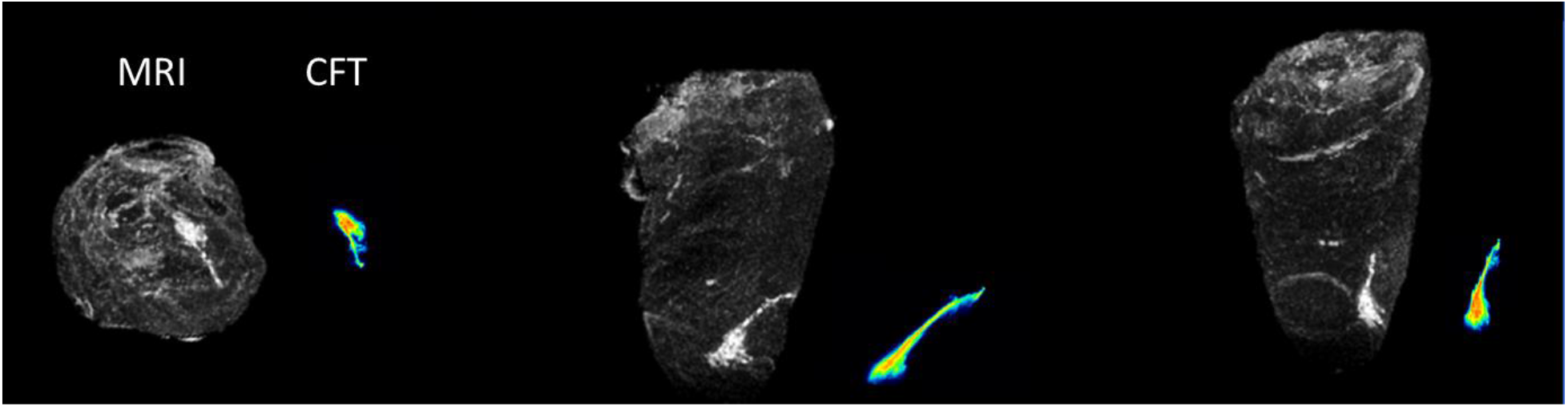
Second example of MRI-based cell tracking of Gd-EOB-DTPA labeled OATP-expressing HEK293 cells transplanted into chicken hearts. Maximum intensity projections from MRI in 3 orthogonal views with corresponding maximum intensity projections from CFT showing corroborating positions.

## Discussion

Hepatic OATPs are poised to make a major impact for molecular imaging by MRI due to their ability to transport FDA-approved MRI contrast agents into cells that express the transporters. We add to this growing field by demonstrating that OATP1B3-expressing cells can be pre-labeled by incubation with Gd-EOB-DTPA prior to injection, and their location post-injection can be readily detected as hyperintense signal by clinically familiar T1-weighted MRI.

One potential advantage of this method for cell labeling and MRI-based cell tracking includes the use of T1-weighted MRI to generate hyperintense signals, rather than T2/T2*-weighted MRI to generate anatomy-obscuring dark contrast. Another potential advantage of this paradigm is the likelihood of contrast agent clearance from the transplantation area in the case of cell death. We hypothesize this to be the case as the small Gd-EOB-DTPA molecule would quickly diffuse and other cells would not accumulate it, even macrophages, due to the lack of OATP transporters. Indeed, the bystander effect from macrophage accumulation of iron oxide nanoparticles from dead cells is a major confounder for MRI-based cell tracking using iron oxide particles^26^. Lastly, this method is unlikely to impact the later re-location of transplanted cells by the established method of IV injection of Gd-EOB-DTPA or Gd-BOPTA as studies have shown that the agent clears from cells between 1 and 5 days^6-12^.

## Conclusion

MRI-based cell tracking of OATP-expressing cells following transplantation is robust and feasible by pre-labeling cells with Gd-EOB-DTPA.

